# Unbiased Classification of the Human Brain Proteome Resolves Distinct Clinical and Pathophysiological Subtypes of Cognitive Impairment

**DOI:** 10.1101/2022.07.22.501017

**Authors:** Lenora Higginbotham, E. Kathleen Carter, Eric B. Dammer, Rafi U. Haque, Erik C.B. Johnson, Duc M. Duong, Luming Yin, Philip L. De Jager, David A. Bennett, James J. Lah, Allan I. Levey, Nicholas T. Seyfried

## Abstract

The hallmark amyloid-β and tau deposition of Alzheimer’s disease (AD) represents only a fraction of its diverse pathophysiology. Molecular subtyping using large-scale -omic strategies can help resolve this biological heterogeneity. Using quantitative mass spectrometry, we measured ~8,000 proteins across >600 dorsolateral prefrontal cortex tissues from Religious Orders Study and Rush Memory and Aging Project participants with clinical diagnoses of no cognitive impairment, mild cognitive impairment (MCI), and AD dementia. Unbiased classification of MCI and AD cases based on individual proteomic profiles resolved three classes with expression differences across numerous cell types and biological ontologies. Two classes displayed molecular signatures atypical of those previously observed in AD neurodegeneration, such as elevated synaptic and decreased inflammatory markers. In one class, these atypical proteomic features were associated with clinical and pathological hallmarks of cognitive resilience. These results promise to better define disease heterogeneity within AD and meaningfully impact its diagnostic and therapeutic precision.

## Introduction

The pathological hallmarks of Alzheimer’s disease (AD), the most common cause of dementia in the elderly [1], include extracellular amyloid-β (Aβ) deposits and intracellular tau neurofibrillary tangles (NFTs) [2, 3]. The long-standing amyloid cascade hypothesis casts Aβ as an early molecular driver of disease, prompting NFT formation, neurodegeneration, and ultimately cognitive decline [4, 5]. While this linear series of pathological events may hold true for rare familial forms of early-onset AD [6], it is now well-recognized that Aβ and tau represent only a fraction of the complex and heterogeneous pathophysiology linked to sporadic late onset AD (LOAD) [7]. Several studies have confirmed that those with AD dementia commonly harbor brain pathologies beyond Aβ plaques and NFTs, including cerebrovascular disease, neocortical Lewy body inclusions, and TAR DNA-binding protein 43 (TDP-43) aggregates [8–12]. Yet, combined with amyloid and tau, these co-pathologies still account for less than half of the variance in cognitive trajectory [13]. Accordingly, genome wide association studies (GWAS) have linked LOAD pathogenesis to a variety of biological mechanisms beyond aberrant protein accumulation and neuronal death, such as glial-mediated inflammation and endothelial integrity [14–21]. These findings highlight the vast pathophysiological heterogeneity underlying cognitive impairment in the elderly.

Molecular subtyping using large-scale -omic strategies promises to resolve this complex biological heterogeneity. Recent genomic clustering of AD based on risk-associated SNP burden revealed disease subgroups linked to distinct biological mechanisms [22]. Subsequent transcriptome-wide studies of the AD brain have identified molecular subtypes corresponding to different combinations of multiple dysregulated pathways [23, 24], including neuroinflammation, synaptic signaling, immune activity, mitochondria organization, and myelination [23]. These studies highlight the utility of large “-omic” datasets in the molecular reclassification of AD and related dementias. Further advancements in such subtyping approaches promise to not only impact diagnostic guidelines, but also enhance the precision of clinical trial recruitment, prognostication, and therapeutic targeting.

To date, large-scale molecular subtyping of AD has primarily focused on genomic and transcriptomic profiles, while protein-based classification remains in its infancy. Yet, marked spatial, temporal, and quantitative differences between mRNA and protein expression make proteomic subtyping a potential source of unique biological insights [25, 26]. Furthermore, compared to RNA differences, protein changes associate more strongly with AD clinical and pathological phenotypes, consistent with their being more proximate mediators of disease manifestations [27–29]. Using unbiased co-expression network analysis, we have demonstrated that the cortical brain regions of those with pathologically defined early- and late-stage LOAD feature a wide range of altered protein systems not observed in the transcriptome [30–33]. These protein alterations correlate strongly with clinical symptoms, biofluid markers, and pathological traits [30–33]. However, it remains unclear whether these protein levels drive distinct molecular subtypes of AD.

To this end, we performed an unbiased proteomic subtyping analysis of mild cognitive impairment (MCI) and AD brain tissues. All samples were derived from the Religious Orders Study or Rush Memory and Aging Project (ROSMAP) longitudinal cohorts, which feature community-based recruitment strategies designed to ensure heterogenous, “real-world” representations of cognitive impairment within the general population [34–36]. Using tandem mass tag mass spectrometry (TMT-MS), we quantified nearly 8,000 proteins across 610 brain tissues from individuals with clinical diagnoses of no cognitive impairment (NCI), MCI, and AD. Unbiased clustering of the nearly 400 MCI and AD tissues by individual proteomic profiles resolved three major classes of cognitive impairment. Here, we thoroughly examine how these classes differ across cell types and biological ontologies. We highlight how two of the three classes harbor proteomic features atypical of molecular trends previously observed in AD neurodegeneration. We also explore how these divergent molecular signatures impact genetic risk and clinicopathologic phenotypes. In sum, our results underscore the biological heterogeneity present among elderly individuals with cognitive impairment and how this translates into differences in cognitive resilience, pathological burden, and genetic risk. Further investigation of these distinct disease subtypes promises to meaningfully impact diagnostic, prognostic, and therapeutic precision in AD.

## Results

### Unbiased classification of the human brain proteome yields three distinct classes of cognitively impaired individuals

The main objective of this study was to classify the brains of those with MCI and AD dementia based on individual proteomic profiles and examine the molecular, genetic, and clinicopathologic features of the resultant subtypes (**Fig. 1**). Using multiplex tandem mass tag mass spectrometry (TMT-MS), we analyzed a total of 610 postmortem dorsolateral prefrontal cortex (DLPFC) tissues from 604 unique individuals enrolled in the Religious Orders Study or Rush Memory and Aging Project (ROSMAP) (**Fig. 1**). These cohorts recruit older individuals without known dementia from United States religious orders, lay retirement centers, senior and subsidized housing communities, and church groups. These participants are then followed longitudinally with cognitive batteries, biospecimen collection, and finally brain autopsy [34–36]. The community-based procedures of ROSMAP were designed to ensure a heterogenous, “real-world” representation of the dementia population found outside of tertiary care centers. Accordingly, clinical diagnoses of NCI, MCI, AD dementia, or other dementia were determined by study experts based principally on clinical history and detailed neuropsychological evaluation [37]. These study procedures have generated cohorts with well-described clinical and pathological heterogeneity, including among participants who ultimately meet neuropathological criteria for AD [12, 35]. To preserve this authentic heterogeneity, we included cases based principally on their clinical consensus cognitive diagnosis (cogdx), a final clinical diagnosis imparted at death by study physicians blinded to neuropathological results. All clinical diagnoses of AD met criteria for possible or probable AD based on National Institute of Neurological and Communicative Disorders and Stroke and Alzheimer’s Disease and Related Disorders Association (NINCDS-ADRDA) guidelines [37, 38].

**Figure 1.**
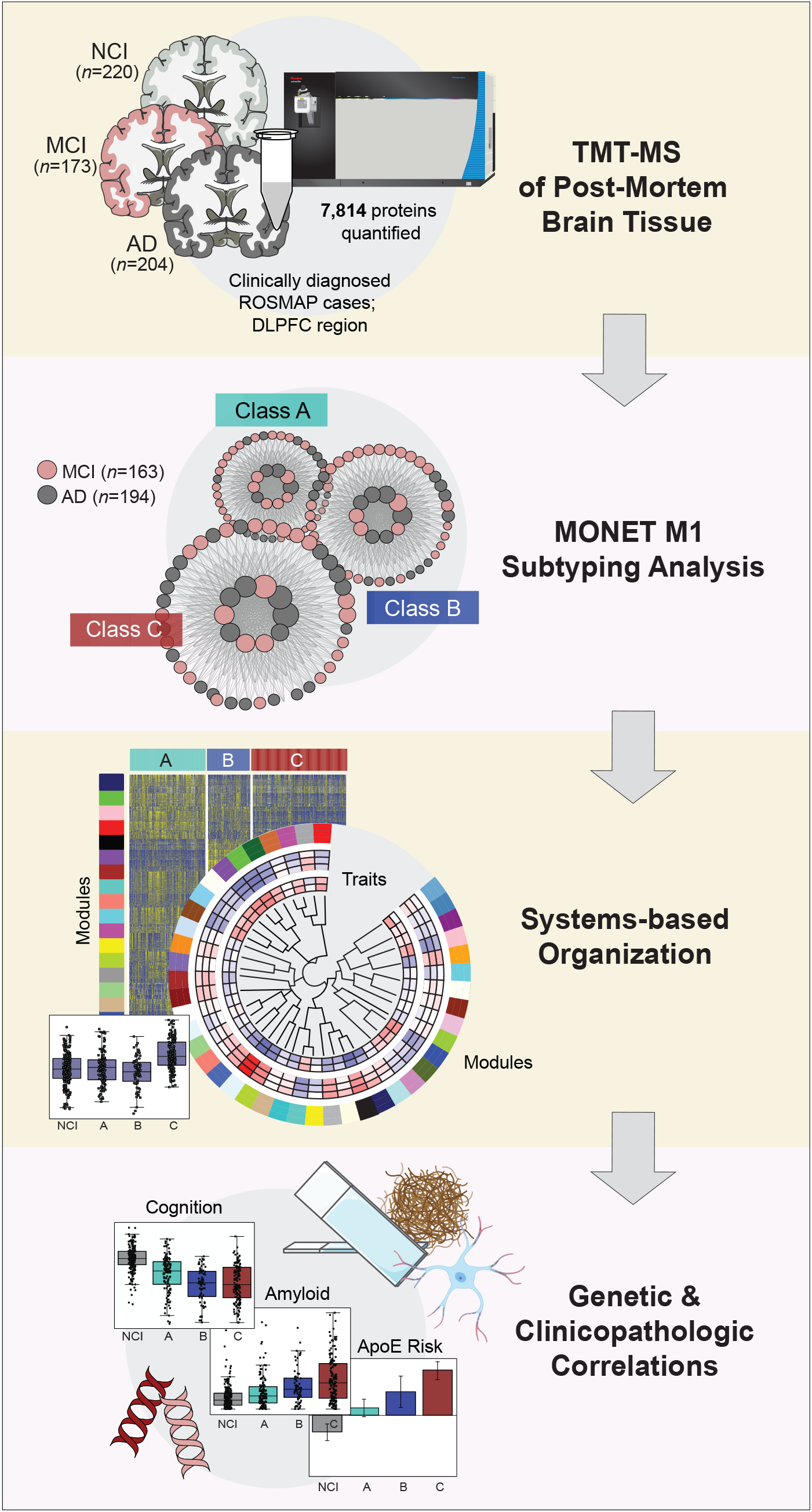
Study Approach. This study classified the brain-derived proteomic profiles of cognitively impaired individuals and characterized these distinct classes using a systems-based biological framework. We first used TMT-MS to analyze >600 DLPFC tissues from ROSMAP participants with clinical diagnoses of NCI (*n*=220), MCI (*n*=173), and AD dementia (*n*=204). We then applied the clustering algorithm MONET M1 to the MCI and AD cases, resolving three proteomic classes termed A, B, and C. We explored biological differences between classes by applying a systems-based organization to these proteomic profiles that was informed by prior network analyses of the AD brain. Finally, we examined genetic, clinical, and pathological differences between NCI and the three classes. Abbreviations: NCI, No Cognitive Impairment; MCI, Mild Cognitive Impairment; AD, Alzheimer’s Disease; TMT-MS, Tandem Mass Tag Mass Spectrometry; DLPFC, Dorsolateral Prefrontal Cortex; ROSMAP, Religious Orders Study and Rush Memory and Aging Project.

TMT-MS quantified 7,814 proteins across 610 ROSMAP tissues with cogdx classifiers of NCI, MCI, or AD dementia. Outlier removal resulted in 597 tissues for subsequent analysis, including 220 NCI, 173 MCI, and 204 AD cases. After adjustments for age, sex, post-mortem interval (PMI), and batch, we clustered the 377 MCI and AD cases into proteomic classes using the statistical algorithm MONET M1 [39]. MONET M1 offers an innovative graph theory approach to module clustering, distinguishing it from more traditional hierarchical algorithms [39, 40]. The parameters of MONET M1 were optimized using a grid search (**Fig. S1**, **Table S1**) to minimize the percentage of cases not assigned to a unique class. Ultimately, 95% (*n*=357) of the 377 cases were assigned to one of three classes, termed A, B, and C (**Fig. 1, Table S2**). Class A (*n*=128) comprised 80 MCI (62%) and 48 AD (38%) cases. Class B (*n*=71) harbored 27 MCI (38%) and 44 AD (62%) cases. Finally, Class C (*n*=158), the largest group, contained 56 MCI (35%) and 102 AD (65%) cases. Given each class comprised a mixture of both MCI and AD cases, we immediately concluded symptom severity was not the only driver of class structure. There were no significant differences in the average age and sex of each class (**Table S3**).

To assess reproducibility of these classes, we employed a bootstrap approach to repeatedly cluster the samples an additional 100 times (**Fig. S2A-B**). On each of these iterations, we applied MONET M1 to a randomly selected 80% (*n*~300) of MCI and AD cases. The resultant clusters generated in each bootstrap iteration were analogous to the original clustering as assessed by strong levels of overlapping class-specific samples (**Fig. S2C**) and highly preserved protein signatures (**Fig. S2D**). Thus, our unbiased classification was highly reproducible, supporting the robustness of MONET M1 in defining consistent patterns of protein expression across cases.

We then independently validated these proteomic classes by applying a second high-performance clustering algorithm to our ROSMAP dataset termed Uniform Manifold Approximation and Projection (UMAP). In recent studies, UMAP has proven capable of effectively reinforcing sample heterogeneity within bulk -omic datasets with clustering structures that maintain biological and clinical meaning [41]. Furthermore, its nonlinear dimension reduction technique has demonstrated meaningful clustering advantages when visualizing high dimensional data compared to traditional linear approaches, such as principal component analysis (PCA) and multidimensional scaling (MDS) [41, 42]. We employed a supervised approach to our UMAP analysis, specifying an output of three distinct clusters. This independent clustering analysis generated three proteomic groups nearly identical to those formed by MONET M1, reinforcing the structure of the original classes (**Fig. S3**). Only one of the 357 cases clustered differently between the algorithms, segregating into Class B with MONET M1 and Class C with UMAP. These results further supported the validity of our three proteomic classes of cognitive impairment.

### Classes differ across a diverse range of disease-associated biological ontologies

We previously showed that the AD cortex features a network of highly reproducible groups or “modules” of co-expressed proteins that reflect disease-associated alterations in a wide range of cell types and molecular functions [30–33]. These network analyses have established a biological framework for the AD brain proteome and its diverse pathophysiology. To provide such biological context to the three classes, we organized their proteomic profiles by the 44 coexpression modules of our deepest AD consensus network, derived from hundreds of tissues in the early and late stages of disease [32]. Approximately 68% of the nearly 8,000 proteins identified among our ROSMAP cases (*n*=5290) mapped to one of these 44 modules. This mirrored previous analyses, which have shown that ~70% of proteins in any given dataset segregate into modules [30–33, 43–48]. The resultant heat map highlights module expression across the three proteomic classes (**Fig. 2A**). Many modules with distinct alterations across classes demonstrated strong associations to specific molecular functions (**Fig. 2A**), cell types (**Fig. 2B**), and clinicopathological traits (**Fig. 2B-C**). See **Table S4** for module correlations to all traits provided for our ROSMAP cohort. Overall, these results showcased differences between classes across a diverse range of disease-relevant biological systems.

**Figure 2.**
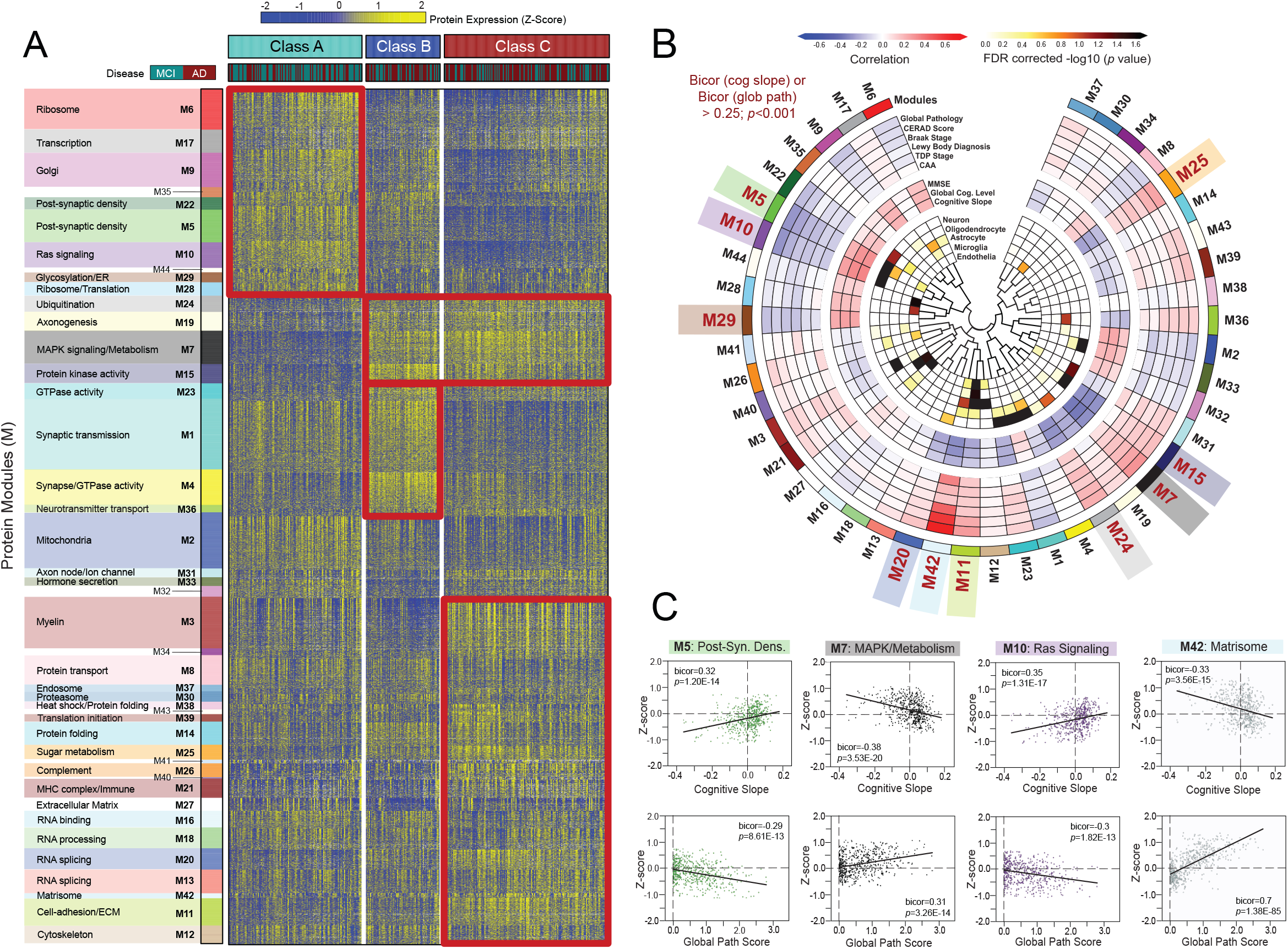
MONET M1 yields three disease-relevant proteomic classes of cognitive impairment. (**A**) Heat map of protein expression across the three proteomic classes generated by MONET M1 analysis. Classes were termed A (*n*=128), B (*n*=71), and C (*n*=158) and each featured a mixture of MCI and AD cases. To provide biological context to the proteomic differences across classes, proteins were organized by modules (M) of co-expression informed by prior AD network analyses. Red boxes highlight modules with relatively elevated levels (yellow shading) in select classes. (**B**) Diagram depicting the associations of each module to cell type and ROSMAP clinicopathological traits. Modules bolded in red (*n*=10) demonstrated exceptionally strong correlations to cognitive slope and/or global pathology (bicor>0.25; *p*<0.001). (**C**) Correlation plots of module abundance (z-score) to cognitive slope or global pathology across all analyzed cases (*n*=610) for select modules with remarkably strong clinicopathological correlations. M5 and M10 demonstrated highly significant positive correlations to cognitive slope and negative correlations to global pathology. In contrast, M7 and M42 were negatively correlated to cognitive slope and positively correlated to global pathology. Bicor correlation coefficients with associated *p* values are shown for each correlation plot. Abbreviations: MCI, Mild Cognitive Impairment; AD, Alzheimer’s Disease; FDR, False Discovery Rate; Post-Syn Dens, Post-Synaptic Density.

Module abundance levels (z-scores) across all cases revealed Class A proteomic signatures most closely matched those of cognitively intact (NCI) individuals, distinguishing this class as the most “control-like” of the three (**Fig. 3A-B, Table S5**). Compared to B and C, Class A featured significantly elevated levels of modules involved in protein synthesis and transport, including M6 (ribosome), M9 (Golgi transport), and M29 (glycosylation / endoplasmic reticulum) (**Fig. 2A and 3A**). Modules linked to RAS signaling (M10) and the post-synaptic density (M5, M22) were also significantly increased in Class A. On the other hand, several modules featured unique decreases in Class A relative to B and C, including those linked to mitogen-activated protein kinase (MAPK) and other kinase-associated pathways (M7, M15) (**Fig. 2A and 3B**).

**Figure 3.**
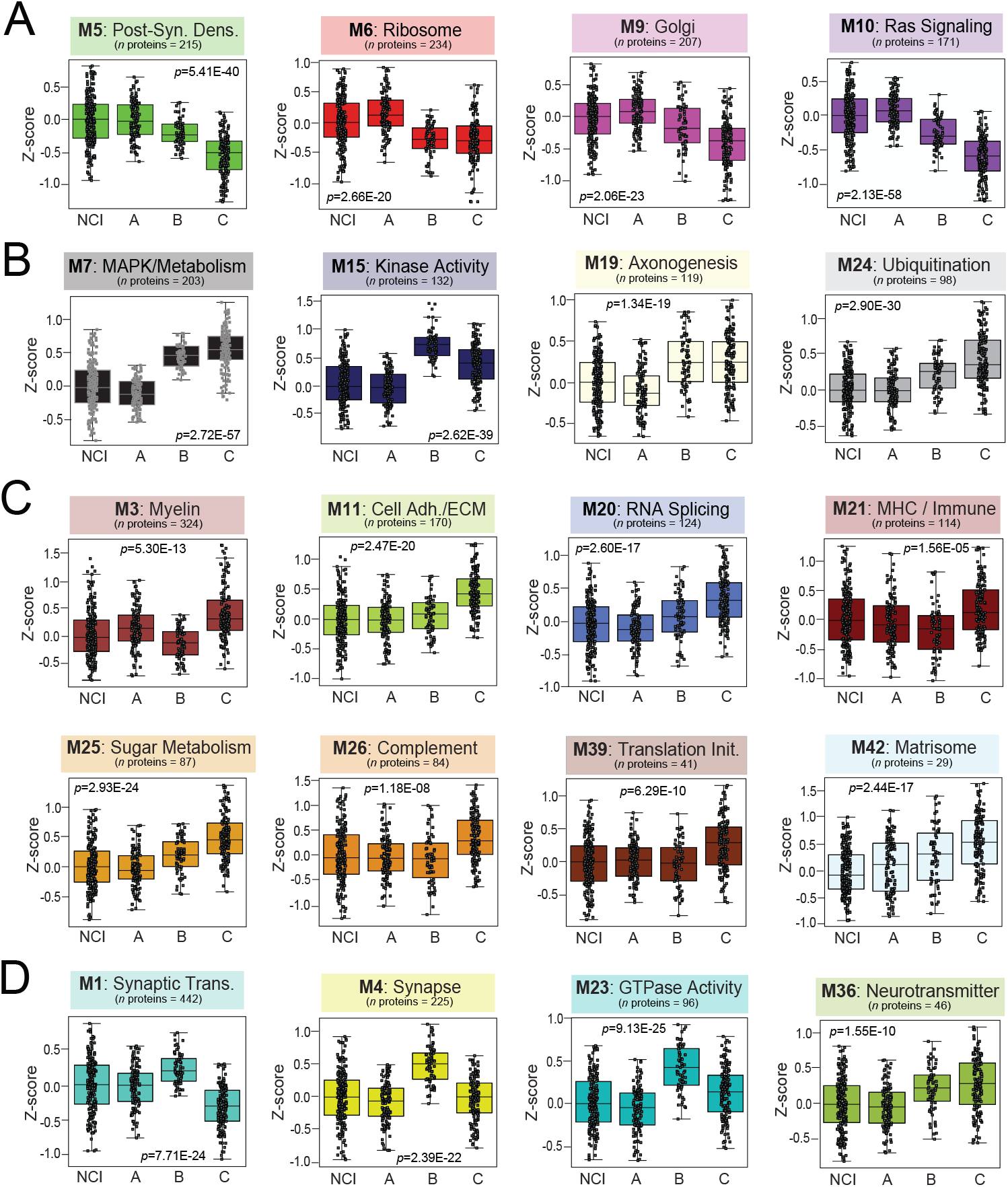
Module abundances highlight class differences across a diverse range of biological ontologies. Abundance levels (z-score) of select modules across NCI cases and the three proteomic classes. ANOVA *p* values are provided for each abundance plot. All modules depicted were significantly altered (*p*<0.001) across the four groups. Box plots represent the median and 25th and 75th percentiles, while box hinges depict the interquartile range of the two middle quartiles within a group. Data points up to 1.5 times the interquartile range from the box hinge define the extent of error bar whiskers. Modules relatively increased in NCI and Class A included M5, M6, M9, and M10, corresponding to post-synaptic density, ribosome, Golgi, and Ras signaling proteins, respectively (**A**). Kinase-associated M7 and M15 were among modules significantly decreased in NCI and Class A compared to the other two classes (**B**). Numerous modules were uniquely increased in Class C, most notably several linked to glial-mediated processes (M3, M11, M21, M26, M42) (**C**). Several synaptic modules (M1, M4, M23, M36) were increased in Class B relative to all other cases (**D**). Abbreviations: NCI, No Cognitive Impairment; MCI, Mild Cognitive Impairment; AD, Alzheimer’s Disease.

Class C featured proteomic changes most consistent with the neurodegenerative trends we have previously observed in pathologically defined AD [31, 32, 45]. Compared to A and B, Class C demonstrated significantly elevated levels of numerous glial-mediated modules linked to inflammation (M26), immune function (M3, M21), and the extracellular matrix (M11, M42) (**Fig. 2A and 3C**). Class C was also the only class that demonstrated significant decreases in M1, a large module linked to synaptic transmission that is consistently depleted in the AD brain (**Fig. 3D**) [30–33, 45]. In contrast, Class B was distinguished from A and C by significant elevations in M1 and several other neuronal modules, including M4 (synapse / GTPase activity), M36 (neurotransmitter transport), and M23 (GTPase activity) (**Fig. 2A and 3D**). These modules were largely associated with the pre-synaptic region and its associated functions (**Fig. 3D**). Meanwhile, post-synaptic modules (M5, M22) were significantly decreased in Classes B and C and remained relatively preserved in Class A. Collectively, these results revealed that in this heterogenous, clinically diagnosed cohort, the proteomic profiles of two-thirds of cognitively impaired cases diverged in key respects from the typical degenerative proteomic changes we have observed previously in pathologically defined tissues.

### Individual protein signatures distinguish classes with high sensitivity and specificity

To identify individual proteins that best discriminate the three classes, we first performed pairwise differential expression analyses. **Figure 4** depicts these volcano plots with individual proteins colored by module membership. As expected, Classes A and C diverged the most with 3,251 significantly altered proteins (*p*<0.05) between them (**Fig. 4A, Table S6**). The “control-like” Class A featured higher levels of several M5 post-synaptic markers (VGF, SYT12, NPTX2), M10 RAS signaling molecules (RASGRF1, ARFGAP2), and M6 mitochondrial ribosome proteins (DAP3, MRPS7, MRPS9, MRPS33, MRPS34). On the other hand, Class C featured increases in numerous proteins from kinase-oriented modules (M7, M15), including MAP kinases (MAPK1, MAP2K6, MAPK3), ribosomal kinases (RPS6KA5), and diacylglycerol kinases (DGKG) (**Fig. 4A**). Large-fold increases in proteins linked to sugar metabolism (M25) and the extracellular matrix (M42) also distinguished Class C from A. These included several highly conserved M42 hubs (SMOC1, MDK, NTN1) repeatedly linked to amyloid burden and APOE-associated risk in prior studies [31, 32]. Meanwhile, Class B pairwise analyses (**Fig. 4B-C**) underscored its unique elevations in neuronal proteins. Several M1 and M4 members (SYN2, NPTXR, SYT17, SYNPR) were significantly increased in Class B compared to A and C.

**Figure 4.**
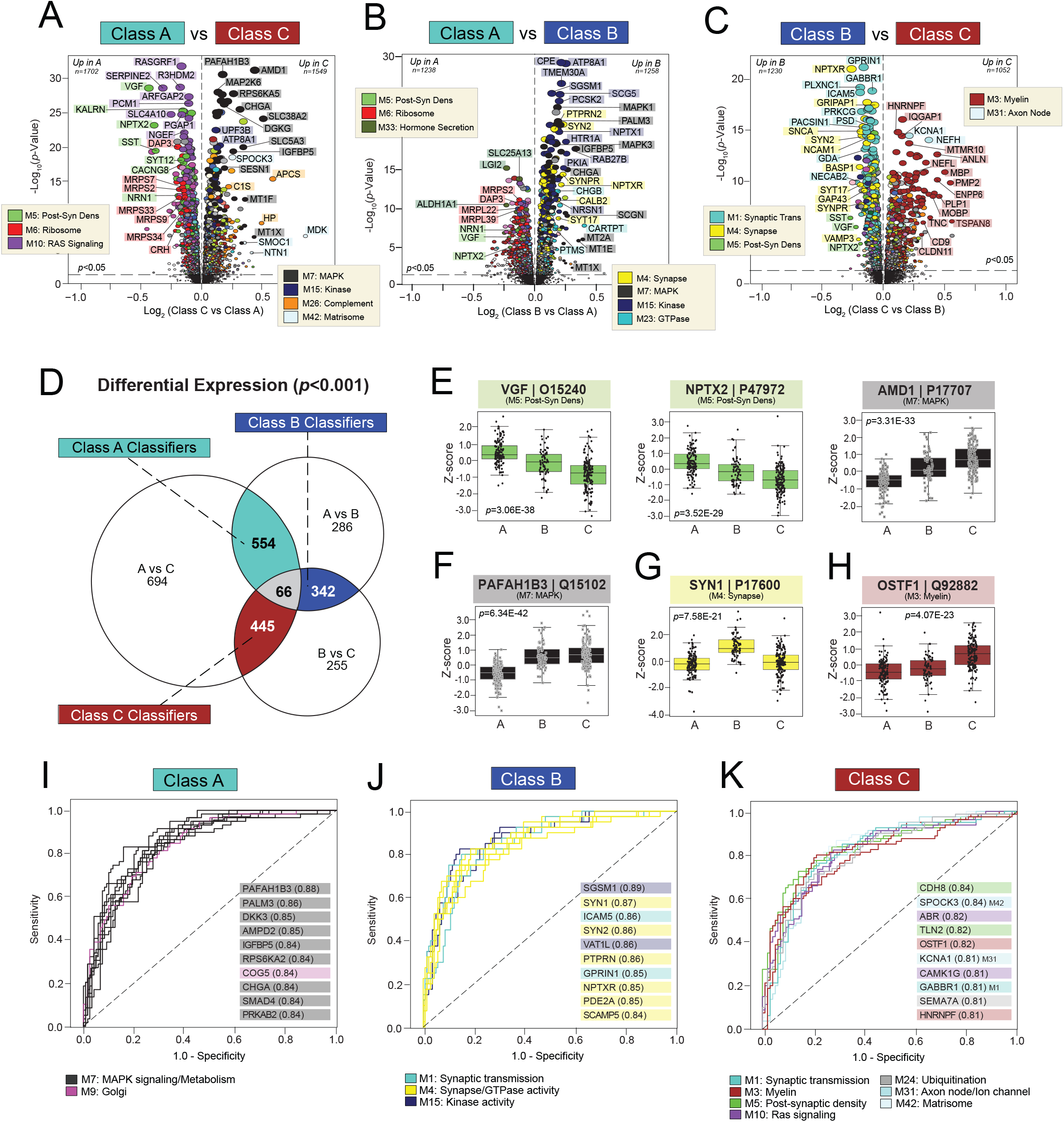
Differential expression of individual proteins reveals highly sensitive and specific classifiers. **A-C**) Volcano plots displaying the log_2_ fold change (x-axis) against the −log_10_ statistical *p* value (y-axis) for proteins differentially expressed between pairwise class comparisons. All *p* values across pairwise comparisons were derived by ANOVA with Bonferroni post-hoc correction. Proteins are shaded according to color of module membership. (**D**) Venn diagram of significantly altered proteins (*p*<0.001) across pairwise class comparisons. There were 66 proteins with significant changes across all three pairwise comparisons, while hundreds of proteins were significantly altered across two of the three pairwise comparisons. The latter were deemed “classifiers”, as each was uniquely altered in one class relative to the other two. There were 554 Class A classifiers, 342 Class B classifiers, and 445 Class C classifiers. (**E-H**) Abundance levels (z-score) of select proteins across NCI cases and the three classes. ANOVA *p* values are provided for each abundance plot. Box plots represent the median and 25th and 75th percentiles, while box hinges depict the interquartile range of the two middle quartiles within a group. Data points up to 1.5 times the interquartile range from the box hinge define the extent of error bar whiskers. The 66 proteins altered across all pairwise class comparisons included neuroprotective markers with well-described links to AD (VGF, NPTX2) and those without known associations to disease (AMD1) (**E**). Classifiers altered in two of the three pairwise class comparisons included PAFAH1B3 for Class A, SYN1 for Class B, and OSTF1 for Class C (**F-H**). (**I-K**) ROC curves of the 10 most sensitive and specific proteins for each class by AUC values, which are included in parentheses. Proteins are shaded according to color of module membership. Abbreviations: Post-Syn Dens, Post-Synaptic Density.

A Venn diagram of significantly altered markers (*p*<0.001) across pairwise class comparisons revealed 66 proteins with significant changes across all three comparisons (**Fig. 4D, Table S7**). These 66 markers included VGF nerve growth factor inducible (VGF), whose levels dropped significantly from Class A to B and then again from B to C (**Fig. 4E**). This M5 neuronal protein is a well-described neuroprotective biomarker with decreased expression in AD brain tissues [49, 50]. Evidence suggests homeostatic VGF signaling promotes cognitive stability, neurogenesis, and synaptic plasticity [49–54]. Thus, its abundance trends in the current study suggested declining neuropreservation from Class A to B to C. Neuronal pentraxin 2 (NPTX2), another contributor to synaptic plasticity with diminished levels in the AD brain [55–57], was also among the 66 markers significantly altered between all three classes. Like VGF, NPTX2 featured steep declines from Classes A to B and B to C (**Fig. 4E**). Trends in these well-described neuroprotective markers underscored the robust proteomic hallmarks of neuropreservation in Class A. In contrast, there were many proteins among these 66 targets that had no known links to AD or neurodegeneration, such as adenosylmethionine decarboxylase 1 (AMD1) which significantly increased from Class A to B to C (**Fig. 4E**).

Our Venn diagram also revealed hundreds of markers significantly altered (*p*<0.001) across two of the three pairwise class comparisons (**Fig. 4D, Table S7**). We referred to these proteins as “classifiers”, as each was uniquely altered in one class relative to the other two. Class A featured 554 classifiers, including platelet activating factor acetylhydrolase 1b catalytic subunit 3 (PAFAH1B3) which displayed markedly decreased levels in Class A relative to B and C (**Fig. 4F**). Decreases in PAFAH1B3 and other members of the kinase associated M7 and M15 comprised nearly 30% of the Class A classifiers. The remaining signatures prominently reflected the increased ribosome (M6), Golgi (M9), and Ras signaling (M10) molecules. Class B was distinguished by 342 classifiers that largely represented increases in several pre-synaptic modules (M1, M4, M23), such as synapsin (SYN1) (**Fig. 4G**). This neuronal protein associates closely with synaptic vesicles and plays a critical role in synaptogenesis and axon development [58]. Finally, Class C classifiers comprised 445 proteins that strongly reflected increases in proteins linked to myelin (M3) and the extracellular matrix (M11), such as osteoclast stimulating factor 1 (OSTF1) (**Fig. 4H**). Decreases in pre- and post-synaptic proteins (M1, M5) were also prominently featured among these Class C signatures.

We assessed the strength of these classifiers by plotting the individual receiver operating characteristic (ROC) curve for each signature in relationship to its associated class. Each curve represented a graphical plot of the true positive rate (sensitivity) against the false positive rate (1-specificity) at various threshold settings. The resultant area under the curve (AUC), a measure of overall classifier performance between values 0 and 1, was then used to identify the strongest signatures for each class (**Fig. 4I-K**, **Table S7**). Kinase-associated proteins (e.g., PAFAH1B3, PALM3, DKK3) were among those most sensitive and specific for Class A (**Fig. 4I**), while Class B was best distinguished by synaptic classifiers (e.g., SYN1, SYN2, GPRIN1, NPTXR) (**Fig. 4J**). Proteins with the highest AUCs for Class C reflected a more diverse set of modules, underscoring the diversity of biological dysregulation in this group. Yet, these Class C signatures still highlighted its prominent synaptic and myelin dysfunction (e.g., CDH8, TLN2, OSTF1, GABBR1, HNRNPF) (**Fig. 4K**). Overall, we concluded that each class featured unique protein signatures capable of distinguishing its members with high sensitivity and specificity.

### Classes demonstrate distinct clinicopathological phenotypes

Given their robust differences in modules linked to clinicopathological traits, we hypothesized our classes would exhibit distinct clinical and pathological phenotypes. To characterize these phenotypic differences, we compared available ROSMAP disease traits directly across classes (**Table S3**). As expected, all three classes demonstrated significant cognitive impairment compared to NCI (**Fig. 5A**). Yet, Class A featured the most preserved cognition among impaired individuals. Class A also displayed the most positive cognitive slopes, indicating a slower rate of decline in these cases (**Fig. 5B**). Accordingly, individual proteins highly correlated to cognitive measures demonstrated starkly different levels in Class A compared to B and C (**Fig. 5A-B**). Post-synaptic markers of M5 (NRN1, NPTX2, OLFM1) were among those most strongly correlated to cognition and displayed precipitous declines from Class A to B and C. Several kinase-associated markers of M7 and M15 (MAP2K6, RPS6KA2, PAFAH1B3, PALM3, TMEM30A) also correlated strongly to cognitive measures but in the opposite direction, with expression levels that sharply increased from Class A to B and C. **Table S8** provides a complete list of proteins significantly correlated to each trait provided for our ROSMAP cohort.

**Figure 5.**
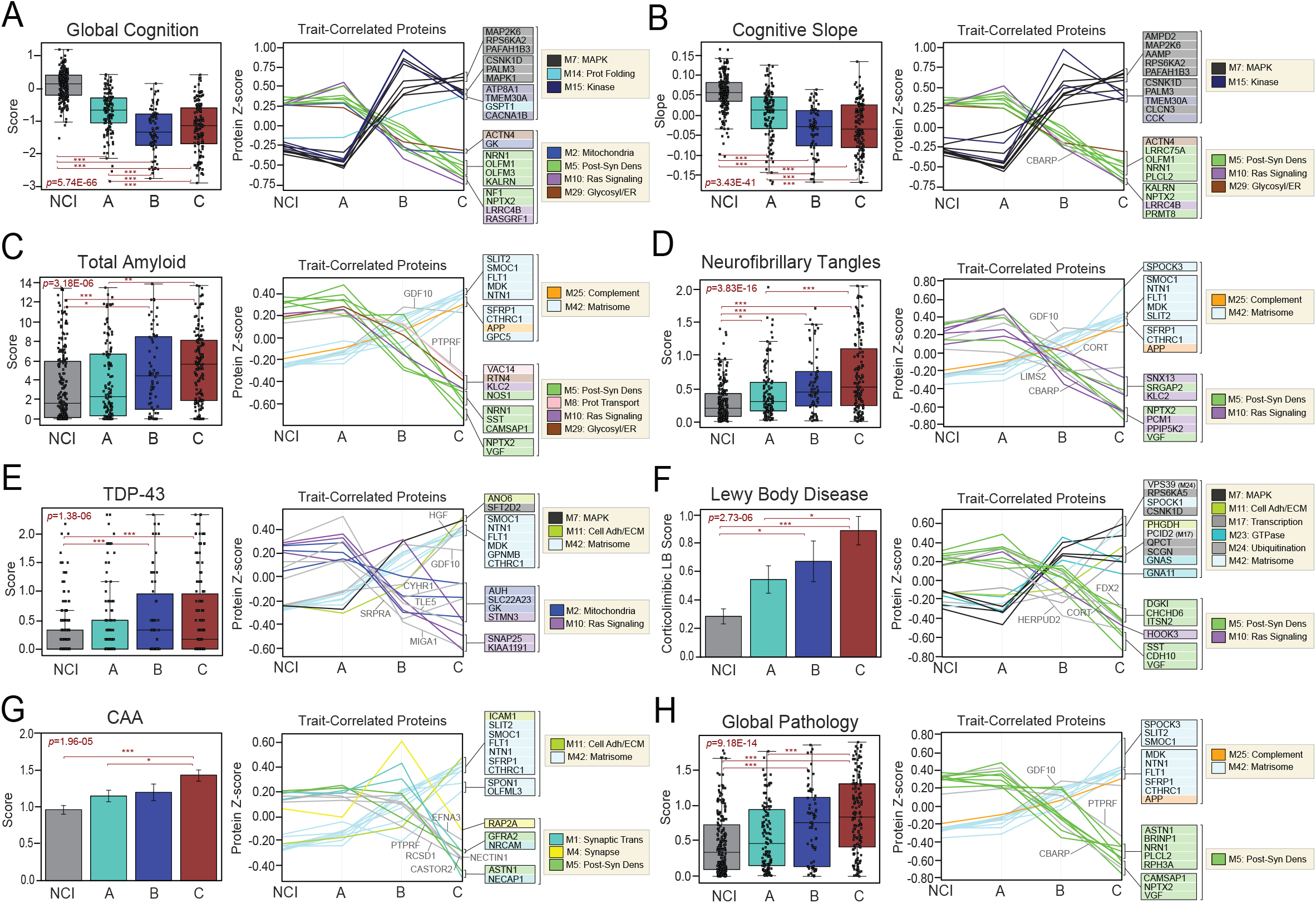
Classes demonstrate different cognitive and pathological features. Cognitive (**A-B**) and neuropathological (**C-H**) characteristics were compared across NCI cases and the three proteomic classes. For each trait, two plots are provided. The first depicts the average scores of each trait across the four groups. The ANOVA *p* value across groups is provided with asterisks indicating statistically significant Tukey post hoc pairwise comparisons (*, *p*<0.05; **, *p*<0.01; ***, *p*<0.001). Box plots represent the median and 25th and 75th percentiles, while box hinges depict the interquartile range of the two middle quartiles within a group. Data points up to 1.5 times the interquartile range from the box hinge define the extent of error bar whiskers. The second plot in each panel showcases the abundance levels (z-scores) across groups of individual proteins highly correlated to that particular trait. The z-scores of the top 10 positively trait-correlated and top 10 negatively trait-correlated proteins are shown. Proteins are shaded according to color of module membership. Proteins without a module assignment are not shaded. Abbreviations: Prot Folding, Protein Folding; Post-Syn Dens, Post-synaptic Density; Glycosyl, Glycosylation; ER, Endoplasmic Reticulum; Prot Transport, Protein Transport; Adh, Adhesion; ECM, Extracellular Matrix.

ROSMAP offers detailed pathological scoring of brain tissues using a variety of semi-quantitative scales [36]. **Fig. 5C-G** showcases the levels of several pathologies across classes, including amyloid plaques, NFTs, cerebral amyloid angiopathy, TDP-43, and Lewy body inclusions. Global pathology scores were also plotted across classes in **Fig. 5H**. As expected, all cognitively impaired cases maintained higher levels of neuropathology compared to NCI. Yet, Class A featured the smallest neuropathological burden of the three subtypes. Its levels of amyloid, tau, and several other pathological features were notably decreased compared to Classes B and C. This was reflected in the global pathology scores of Class A, which were the lowest of all cognitively impaired cases. In contrast, Class C featured the highest burden of global pathology among the three subtypes, surpassing A and B in average levels of NFTs, Lewy body inclusions, and CAA. Proteins highly correlated to neuropathological measures were generally most altered in Class C compared to all other groups (**Fig. 5C-H**, **Table S8**). Post-synaptic (M5) and matrisome (M42) markers were most consistently reflected among pathology-associated markers. Accordingly, global pathology trends were most strongly correlated to post-synaptic (e.g., NRN1, NPTX2, RPH3A, VGF) and matrisome markers (e.g., SPOCK3, SMOC1, MDK, NTN1, FLT1) (**Fig. 5H**).

Overall, these findings highlighted distinct clinicopathological phenotypes across our proteomic classes. Most notably, these results highlighted greater levels of cognitive stability in Class A, consistent with the robust neuroprotective trends in its proteomic profile. Meanwhile, Class C demonstrated the highest levels of neuropathology, aligning with its prominent neurodegenerative proteomic signatures.

### Class C proteomic signatures strongly mirror those of high-risk ApoE4 carriers

Polymorphic alleles in the *APOE* gene are the strongest known genetic determinants of LOAD risk [59–61]. Individuals carrying the E4 allele are at increased risk for AD development compared to those with the more common E3 allele. Meanwhile, a copy of the E2 allele is neuroprotective and decreases the risk of LOAD. We have previously demonstrated that *APOE* genotype and its associated risk strongly correlate with a variety of protein modules in the human AD brain, spanning metabolism, inflammation, synaptic activity, and other molecular functions [43]. We have also shown that the expression patterns of certain modules, such as the matrisome-associated M42, are genetically regulated by the *APOE* locus [32]. Therefore, we hypothesized that our proteomic classes would differ in levels of *APOE*-related risk and associated protein signatures.

Analysis of genotype composition across classes revealed a mixture of high-risk (E3/4, E4/4), risk-neutral (E3/3), and low-risk (E2/2, E2/3) genotypes in each class. E3/3 was the most abundant genotype present throughout the dataset, comprising 60-70% of cases in each class (**Fig. 6A**). High-risk E4 carriers (E3/4, E4/4) were second most abundant, though less evenly distributed. Class C featured over twice as many E4 carriers (*n*=46, 29%) as Classes A (*n*=21, 16%) and B (*n*=17, 24%). Low-risk E2 carriers (E2/2, E2/3) were notably less abundant than high-risk cases, accounting for only 12% (*n*=16) of Class A, 10% (*n*=8) of Class B, and 6% (*n*=10) of Class C. Finally, E2/4 cases were rare and comprised no more than 3% of any class. A comparison of overall *APOE*-associated risk revealed all three classes featured higher risk levels compared to NCI cases. Yet, Class C displayed significantly higher risk compared to Classes A and B (**Fig. 6B**). Collectively, these results suggested that class structure was not solely determined by *APOE* genotype. Yet, the three classes did meaningfully differ in their proportions of high-risk E4 carriers and average levels of genetic risk.

**Figure 6.**
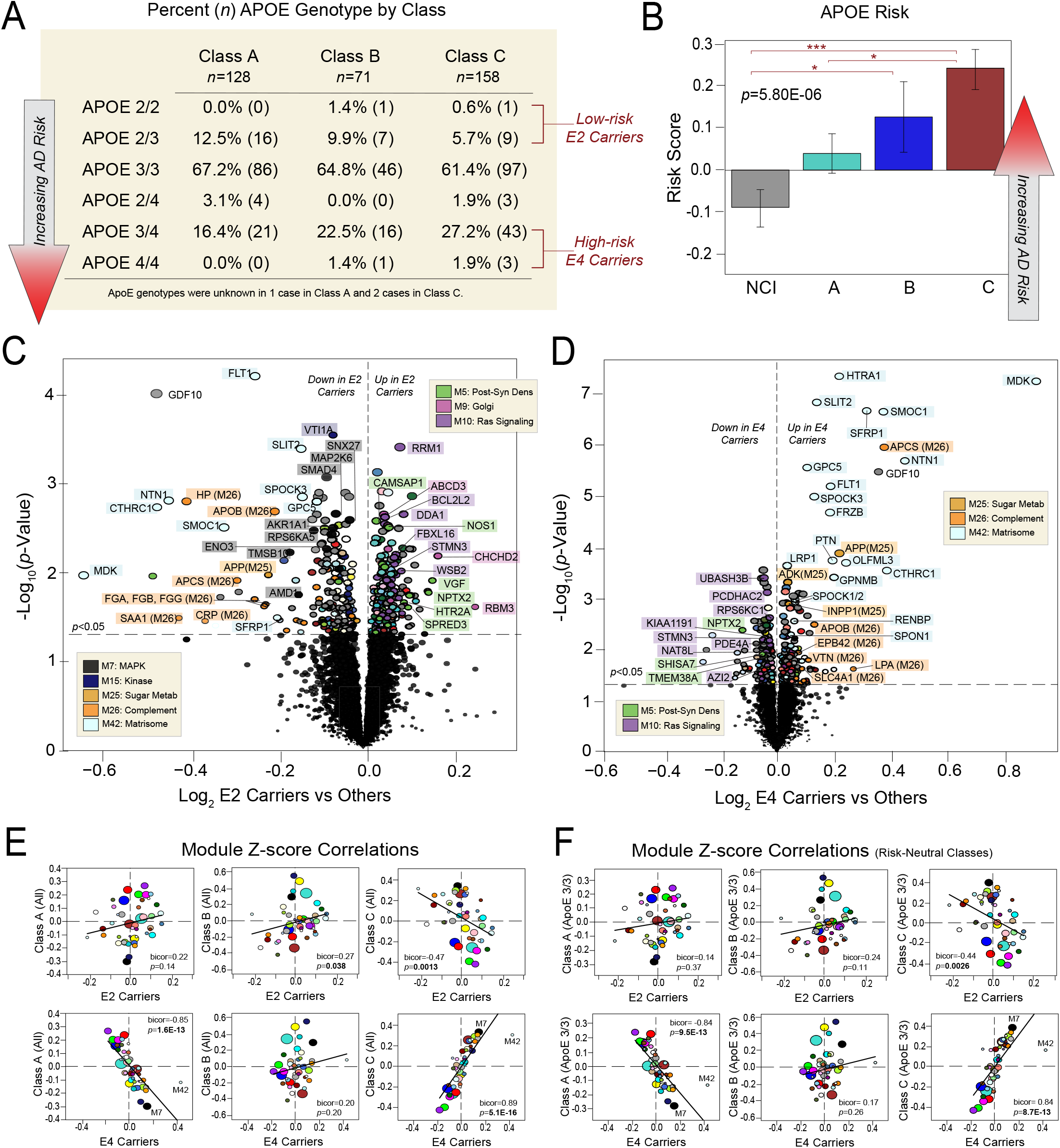
Class C protein expression strongly mirrors that of ApoE4 carriers. (**A**) Table showcasing the percentages of different *APOE* genotypes within each class. The corresponding number of cases with each genotype is also provided in parentheses. Cases considered low-risk E2 carriers or high-risk E4 carriers are indicated. Class C comprised twice as many high-risk E4 carriers compared to Classes A and B. (**B**) Comparison of average *APOE* risk scores across NCI and the three proteomic classes. Individual risk scores for each case were calculated by assigning −1 points to each E2 allele, 0 points to each E3 allele, and +1 points to each E4 allele. The ANOVA *p* value across groups is provided with asterisks indicating statistically significant Tukey post hoc pairwise comparisons (*, *p*<0.05; **, *p*<0.01; ***, *p*<0.001). (**C-D**) Volcano plots displaying the log_2_ fold change (x-axis) against the *t* test-derived −log_10_ statistical *p* value (y-axis) for proteins differentially expressed in E2 carriers or E4 carriers when compared to all other cases, excepting those with E2/4 genotypes which were excluded from these analyses. Thus, (**C**) is a comparison of protein expression in the 34 cases with E2/2 and E2/3 genotypes to the 313 cases with E3/3, E3/4, and E4/4 genotypes, while (**D**) is a comparison of protein expression in the 84 cases with E3/4 and E4/4 genotypes to the 263 cases with E2/2, E2/3, and E3/3 genotypes. Proteins are shaded according to color of module membership. (**E-F**) Correlation plots of module abundance levels (z-scores) in E2 (E2/2, E2/3) or E4 (E3/4, E4/4) carriers to those of each proteomic class. Class-specific z-scores in (**E**) reflect all members of each class, while those in (**F**) reflect only individuals with E3/3 genotypes in each class. Bicor correlation coefficients with associated *p* values are shown for each correlation plot. Abbreviations: Post-Syn Dens, Post-Synaptic Density; Metab, Metabolism.

We then sought to compare risk-associated protein signatures across classes. First, we identified those protein alterations most strongly linked to *APOE* carrier status, regardless of class (**Table S9**). **Fig. 6C** showcases proteins significantly altered in E2 carriers (E2/2, E2/3) versus other cognitively impaired cases. E2 carriers demonstrated stark decreases in kinase related (M7, M15) proteins and increases in post-synaptic (M5), Golgi (M9), and Ras signaling (M10) markers. In contrast, E4 carriers (E3/4, E4/4) featured decreases in post-synaptic (M5) and Ras signaling (M10) proteins when compared to other cognitively impaired cases (**Fig. 6D**), as well as significant increases in proteins linked to sugar metabolism (M25), immune function (M26), and the matrisome (M42). As expected, hub proteins of M42 (SMOC1, MDK, NTN1) were among those markers most elevated in E4 carriers, consistent with our previous findings that this module is under control of the *APOE* locus [32]. Accordingly, LDL receptor related protein 1 (LRP1), another M42 member and known APOE interactor [62, 63], was also significantly elevated in E4 carriers.

To examine these risk-associated protein signatures across classes, we then correlated the proteomic profiles of E2 and E4 carriers with those of each class. E2-associated module expression was positively correlated to module expression in both Classes A and B (**Fig. 6E, Table S10**). However, only its correlation with Class B reached statistical significance (bicor=0.27, *p*=0.038). E2 module expression also significantly correlated to that of Class C, but in the negative direction (bicor=-0.47, *p*=0.0013). In stark contrast, E4 module expression featured remarkably strong negative correlations to Class A (bicor=-0.85, *p*=1.6E-13) and positive correlations to Class C module expression (bicor=0.89, *p*=5.1E-16) (**Fig. 6E**). E4 expression demonstrated no significant correlation to that of Class B (bicor=0.20, *p*=0.20). To ensure that these results were not driven by a minority of E2 or E4 carriers in each class, we repeated all six correlations using the module expression of only E3/3 cases in each class (**Fig. 6F, Table S10**). These results were nearly identical to those of the initial correlations. Notably, the strong positive association between E4 and Class C module expression was maintained (bicor=0.84, *p*=8.7E-13). Thus, we concluded that irrespective of their individual genotypes, Class C cases harbored proteomic profiles highly similar to those of high-risk E4 carriers. This supported the conclusion that with its heightened inflammatory signatures, steep cognitive slopes, and exceptionally elevated neuropathological burden, Class C reflected a high-risk state of cognitive impairment.

## Discussion

The diagnosis, monitoring, and treatment of AD are currently limited by biomarker tools that fail to capture its vast pathophysiological heterogeneity. Large-scale molecular subtyping promises to resolve this heterogeneity and enhance diagnostic and therapeutic precision in AD. To this end, we used an unbiased proteomic approach to subtype nearly 400 ROSMAP brain tissues from clinically diagnosed MCI and AD cases. We resolved three classes among these cognitively impaired individuals, each driven by proteomic changes across a variety of cell types and biological ontologies. All classes featured a mix of mildly impaired and demented individuals, indicating clusters driven by more than clinical severity at death. Accordingly, further examination of these classes highlighted distinct genetic, clinical, and pathological phenotypes.

Class C featured the most neurodegenerative proteomic profile of the three groups. Synaptic loss and heightened glial activation were among its most distinct proteomic signatures. Glial-enriched modules associated with myelin (M3), immune function (M21), complement pathways (M26), and the matrisome (M42, M11) were distinctly elevated in Class C compared to A and B. These glial signatures strongly correlated to the exceptionally high neuropathological burden of Class C. Hub proteins of M42 and M11 were among those most strongly correlated to neuropathology. We and others have previously linked many of these matrisome markers (e.g., SMOC1, NTN1, MDK) to Aβ accumulation [31–33, 64]. Yet, the current study also showcased their strong associations to various non-amyloid pathologies, including NFT, CAA, and TDP-43 deposition. Accordingly, Class C demonstrated distinctly elevated levels of mixed global neuropathology, extending beyond amyloid and tau accumulation. This high pathological burden aligned with the overall aggressive phenotype of Class C, which also featured steep cognitive slopes and elevated levels of genetic risk. In fact, our results suggested that a Class C proteomic profile was nearly equivalent to that of a high-risk E4 carrier.

In contrast, Class A featured the most neurologically preserved proteomic profile, closely mirroring NCI cases in synaptic, metabolic, and inflammatory signatures. Class A was best distinguished from B and C by decreases in kinase-associated modules (M7, M15) and increases in RAS signaling proteins (M10). RAS signaling molecules are known to regulate various aspects of the MAP kinase (MAPK) pathway [65–67], indicating biologically meaningful links between these Class A signatures. The proteomic hallmarks of Class A correlated strongly to cognitive trajectory. Of all 44 modules, M7 demonstrated the most robust correlations to cognitive slope, with lower protein levels indicating increased cognitive stability. Thus, the cognitive slope of Class A was significantly more stable compared to Classes B and C. Class A, while demonstrating higher amyloid and tau deposition relative to NCI, also displayed the smallest burden of global neuropathology among those with cognitive impairment. In addition, its average *APOE* risk was significantly lower than that of Classes B and C, and its protein expression strongly anti-correlated to that of high-risk E4 carriers. Collectively, these findings showcased the milder, less aggressive disease phenotype of Class A. Accordingly, its most highly sensitive and specific classifiers included various proteins linked to neurologic resilience, such as Ras protein specific guanine nucleotide releasing factor 1 (RASGRF1), an important regulator of neural plasticity with links to hippocampus-dependent memory [68–73]. Also among Class A classifiers was neuritin (NRN1), an M5 synaptic protein that has strongly associated with cognitive resilience in prior proteomic studies [74] and has known roles in synaptic maturation and stability [75–77].

Class B displayed a proteomic profile largely intermediate to the extremes of Classes A and C. Class B demonstrated clear degenerative changes relative to Class A, including increases in kinase modules (M7, M15) and decreases in RAS signaling proteins (M10). Yet, Class B lacked many of the hallmarks of glial activation observed in Class C. The expression of known neuroprotective markers underscored Class B as an intermediate state. Levels of VGF and NPTX2, neuroprotective markers that typically decrease in the degenerating brain [49–57], were highest in Class A and lowest in Class C, leaving B in between. However, the proteomic profile of Class B was not entirely transitional in nature. Class B was distinguished by its markedly elevated levels of select neuronal modules (M1, M4, M23, M36) compared to both Classes A and C. These Class B neuronal signatures strongly reflected pre-synaptic functions, including neurotransmitter transport, GTPase activity, and signal transmission. These neuronal modules did not correlate strongly to any genetic, clinical, or pathological traits. Therefore, it is unclear what function these synaptic signatures serve for Class B and whether they comprise a hallmark of neuronal resilience or dysfunction. In addition, what impact these heightened levels of pre-synaptic proteins have on the marked decreases in post-synaptic modules (M5, M22) also observed in Class B is unclear.

Overall, these results revealed only a third of cognitively impaired individuals with clinical MCI and AD harbor characteristic proteomic signatures of neurodegeneration. The remaining cases displayed atypical molecular signatures, which in some cases were strongly correlated to cognitive resilience. These results align to some extent with recent transcriptomic analyses, which also identified “atypical” RNA profiles in over half of MCI and AD brains [23, 24]. However, several key protein modules differentially expressed across classes are not observed in the AD transcriptomic network [32]. One example is M7, a module strongly linked to cognitive trajectory and whose hubs are strong Class A classifiers. Despite these robust disease associations, this module is not preserved in the AD network transcriptome [32, 33, 45]. Another such module not reflected in the AD transcriptomic network is M42 [32], which demonstrated remarkably strong neuropathologic associations and comprises hubs sensitive and specific for Class C.

Genetic risk did not exclusively dictate proteomic classification, as cases with low- and high-risk *APOE* genotypes were scattered throughout all classes. The proteomic profile of Class C strongly mirrored that of E4 carriers, but this robust association persisted regardless of whether individuals in Class C carried an E4 allele. The strong anti-correlations observed between Class A and E4 module expression were also independent of individual Class A genotypes. Meanwhile, Class B module expression was not correlated at all to that of E4 carriers and only weakly to that of E2 carriers. Of note, it is possible that given our generally low numbers of E2 carriers among cognitively impaired cases (*n*=34), we were simply underpowered to detect more robust correlations to low-risk protein signatures. This would explain why E2 proteomic signatures mirrored several trends observed in Class A (e.g., elevated RAS signaling and post-synaptic proteins) but the two failed to demonstrate statistically significant correlations.

Other limitations of the current study included a lack of racial diversity among analyzed cases. Using a community-based, clinically diagnosed cohort ensured clinical and pathological heterogeneity. Yet, our analyses were limited largely to non-Hispanic white individuals. Thus, it is unclear if the same classes would be detected in a more racially diverse analysis. Growing evidence indicates that cerebrospinal fluid (CSF) tau and other molecular markers of AD require adjustments for race [78, 79], suggesting this variable could significantly impact pathophysiological classification of disease. In addition, because we regressed for age and sex prior to clustering, we have a limited understanding of how these variables might also influence subtyping results. Thus, it will be important for future investigations to examine the effects of these demographic factors on proteomic clustering.

Additional studies examining the representation of these brain-derived classes in the CSF and plasma proteomes will also be critical to clinical translation. Recent studies integrating the AD brain and biofluid proteomes have revealed that many key disease-associated brain modules are highly represented in CSF [30, 31]. In fact, we have shown that alterations in the AD CSF proteome reflect a diverse range of brain-based pathophysiology, including synaptic, vascular, inflammatory, and metabolic dysfunction [30]. Thus, the AD CSF proteome promises to mirror the brain with distinct classes featuring unique protein signatures and clinicopathological phenotypes. A recent subtyping analysis of AD CSF based on the levels of ~700 proteins identified subtypes of disease with distinct molecular signatures [80]. Yet, larger-scale integration studies of the brain and CSF proteomes are required to identify biofluid subtypes that best reflect cortical hallmarks of cognitive resilience and global pathology. Such efforts to refine classes with close links to brain-based pathophysiology will be key to meaningfully advancing diagnostic and therapeutic precision in AD.

## Supplementary Materials

Figure S1: MONET M1 grid search and parameter selection

Figure S2: Validation of MONET M1 results using iterative bootstrapping

Figure S3: UMAP Analysis Reinforces proteomic classes of cognitive impairment

Table S1: Grid Search for MONET M1 Analysis

Table S2: Case kMEs for MONET M1 Analysis

Table S3: Demographics and Clinicopathological Traits across Classes

Table S4: Module Abundance Correlations to Demographics and Clinicopathological Traits

Table S5: Differential Module Abundance across Classes

Table S6: Differential Protein Abundance across Classes

Table S7: Venn Diagram Results and AUC Values of Top Protein Classifiers

Table S8: Protein Abundance Correlations to Demographics and Clinicopathological Traits

Table S9: Differential Protein Abundance across E2 and E4 Carriers

Table S10: Module Abundance across Classes and APOE Subgroups

---

**Figure S1.**
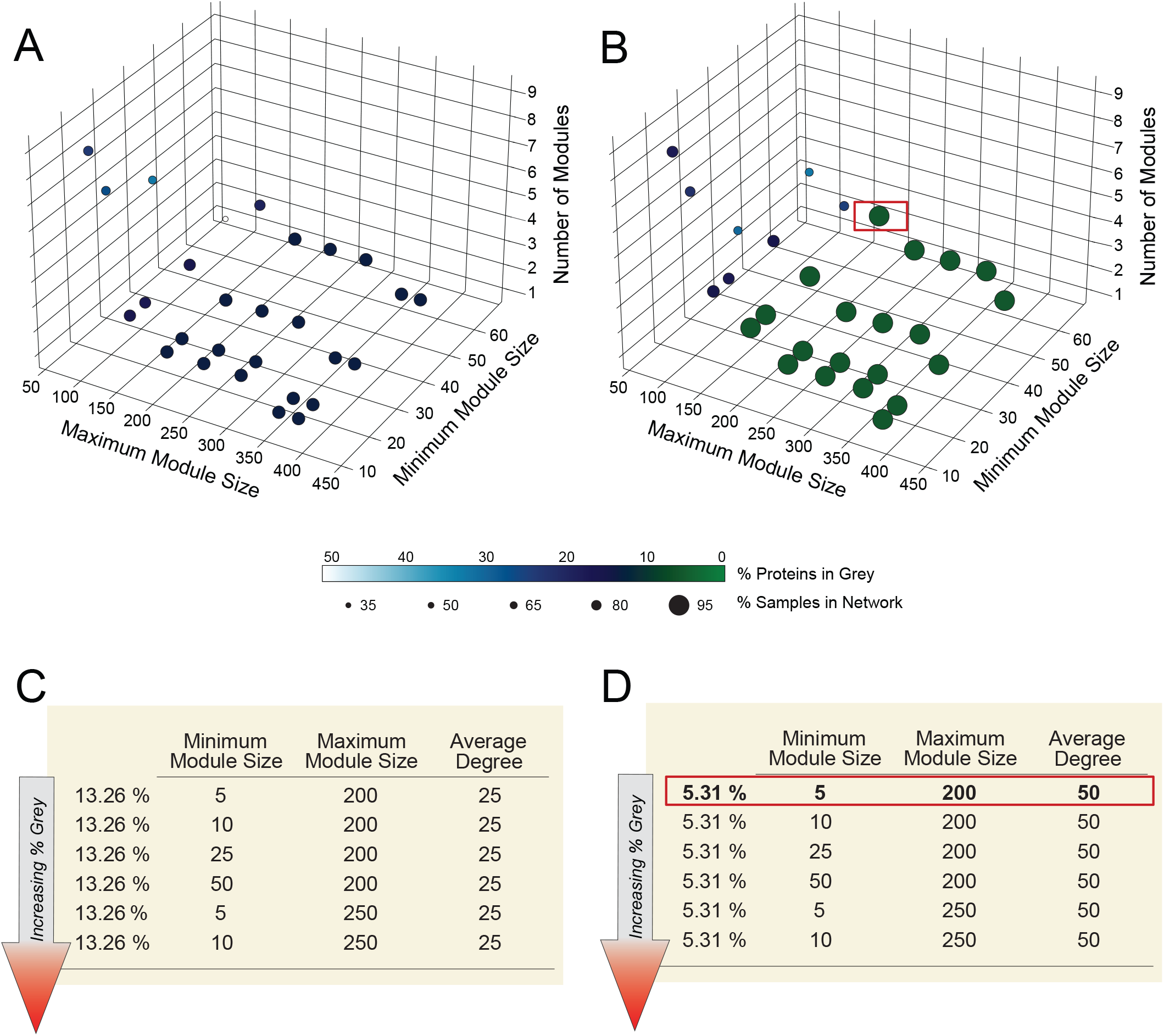
MONET M1 grid search and parameter selection. To select the hyperparameters used in the MONET M1 clustering, a grid search of varying minimum module size, maximum module size, and average degree was performed. The size and percent unassigned cases (grey) for the clusters produced in each combination are visualized as a 3D plot of minimum module size, maximum module size, and number of clusters produced (**A** desired average degree = 25 and **B** desired average degree = 50). The top performing parameter sets as determined by minimal percent grey are shown in panel **C** and **D**. Generally, assigning the desired average degree to 50 decreases the percent grey in each of the tested parameter sets. The hyperparameters selected were minimal module size = 5, maximum module size = 200, desired average degree = 50. The results of these parameters are indicated by red boxes in panel **B** and **D.**

**Figure S2.**
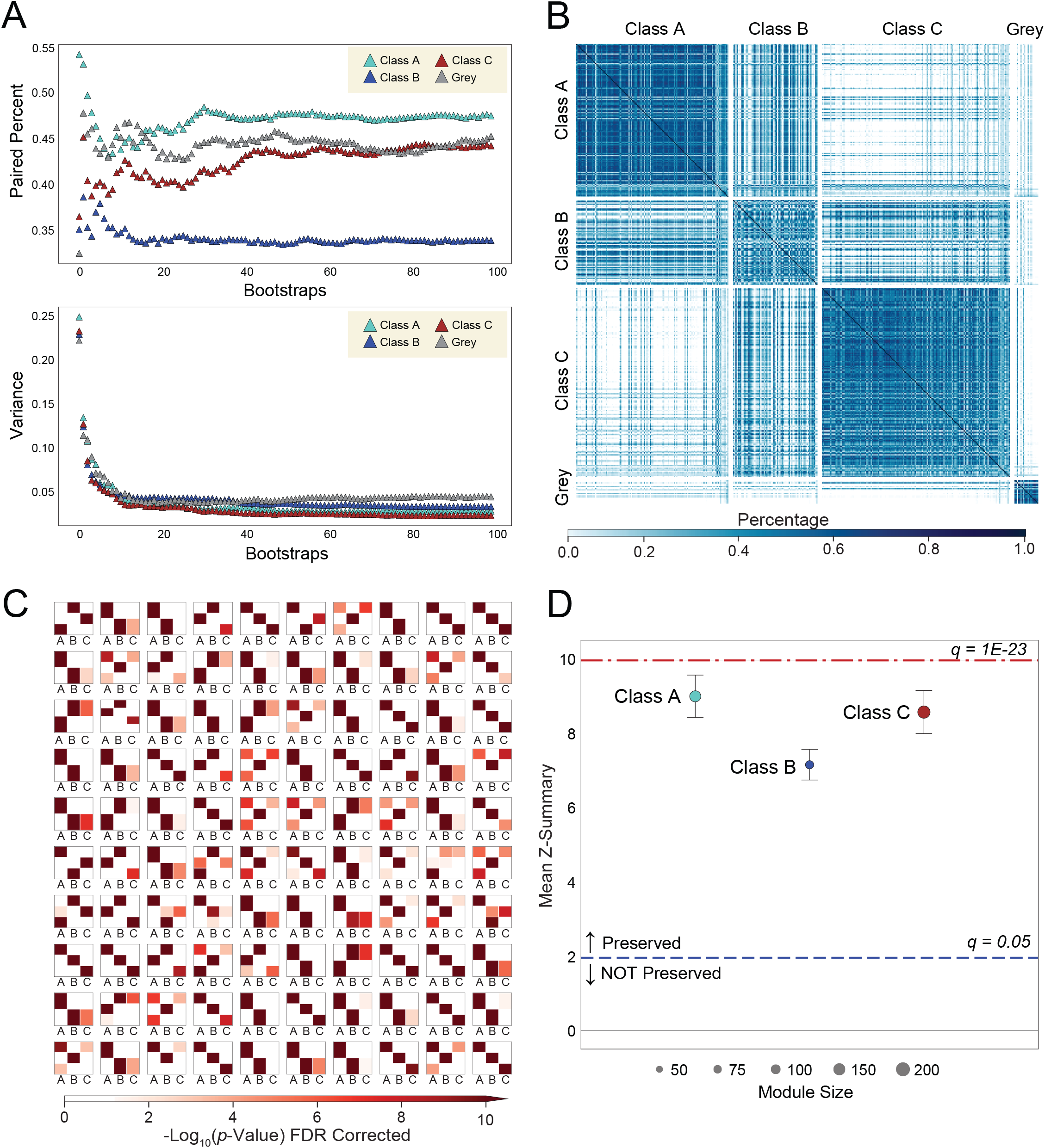
Validation of MONET M1 results using iterative bootstrapping. The rate at which sample pairs are assigned to the same cluster (termed the paired percentage) were calculated for every bootstrapped run (80-20 split). The average paired percentage per cluster and variance of the percentage were tracked to ensure the bootstrapping was well converged (**A**). A heatmap of the pairwise cluster rate shows three distinct clusters (**B**). Hypergeometric Fisher’s exact test (FET) and module preservation was run on each of the MONET M1 reclustering steps with the original MONET M1 network as a reference. FET results revealed significant class-specific sample overlap (FDR-corrected *p*<0.01) across the 100 iterations (**C**). A mean Z_summary_ score was calculated and demonstrates that each of the MONET M1 classes are well preserved (q < 0.05) in each of the 100 bootstrap steps (**D**).

**Figure S3.**
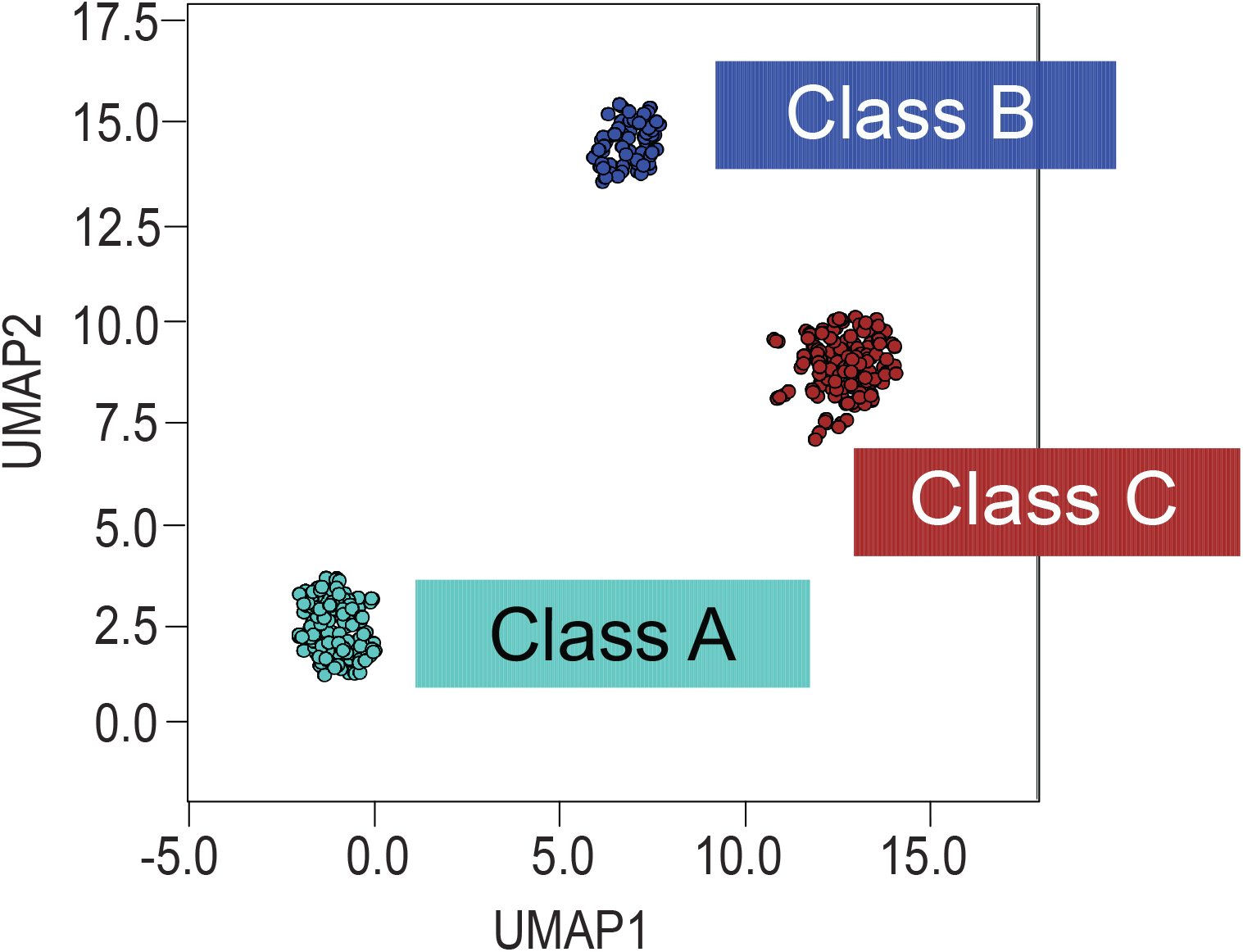
UMAP analysis reinforces proteomic classes of cognitive impairment. Uniform Manifold Approximation and Projection (UMAP) supervised clustering algorithm segregated cognitively impaired ROSMAP cases into three classes nearly identical to those formed by MONET M1. Only one of the 357 cases clustered differently between the algorithms, segregating into Class B with MONET M1 and Class C with UMAP.

## Materials and Methods

### Brain Tissue

DLPFC tissues from Brodmann area 9 (BA9) were obtained from the autopsy collections of the Religious Orders Study or Rush Memory and Aging Project [34–36]. Both studies were approved an Institutional Review Board of Rush University Medical Center. All participants signed an informed consent, an Anatomic Gift Act, and a repository consent allowing their resources to be repurposed with appropriate inter-institutional agreements. ROSMAP features community-based cohorts, which recruit older individuals without known dementia from United States (US) religious orders, lay retirement centers, senior and subsidized housing communities, and church groups. These participants are then followed longitudinally with cognitive batteries, biospecimen collection, and finally brain autopsy [34–36]. All participants are assigned a clinical consensus cognitive diagnosis (cogdx) at death, derived by study experts blinded to postmortem neuropathology. The cogdx scale includes values of 1 (NCI), 2 (MCI and no other cause of cognitive impairment [in addition to AD]), 3 (MCI and another cause of cognitive impairment [in addition to AD]), 4 (AD and no other cause of cognitive impairment), 5 (AD and another cause of cognitive impairment), and 6 (other dementia). All diagnoses of AD met criteria for possible or probable AD based on National Institute of Neurological and Communicative Disorders and Stroke and Alzheimer’s Disease and Related Disorders Association (NINCDS-ADRDA) guidelines [37, 38]. Only cases with cogdx classifiers of 1-5 were included in the current study. Cases with cogdx classifiers of 6 were excluded. We also excluded rare cases with cogdx classifiers that did not align with cognitive battery scores. ROSMAP cases are richly characterized using a variety of clinical and pathological traits that were used to describe the case grouping generated in our clustering analysis [34, 36]. Postmortem neuropathological traits of interest included neuritic plaque distribution, which was scored according to the Consortium to Establish a Registry for Alzheimer’s Disease (CERAD) criteria [81] and extent of neurofibrillary tangle pathology, which was assessed with the Braak staging system [82]. Other neuropathologic diagnoses and clinical traits were made in accordance with established criteria and guidelines [83]. All case metadata are provided in https://www.synapse.org/ADsubtype.

### Brain Tissue Homogenization and Protein Digestion

Tissue homogenization was performed essentially as described [33, 84]. Approximately 100 mg (wet tissue weight) of brain tissue was homogenized in 8 M urea lysis buffer (8 M urea, 10 mM Tris, 100 mM NaH_2_PO_4_, pH 8.5) with HALT protease and phosphatase inhibitor cocktail (ThermoFisher) using a Bullet Blender (NextAdvance). Each Rino sample tube (NextAdvance) was supplemented with ~100 μL of stainless-steel beads (0.9 to 2.0 mm blend, NextAdvance) and 500 μL of lysis buffer. Tissues were added immediately after excision and homogenized with bullet blender at 4 °C with 2 full 5 min cycles. The lysates were transferred to new Eppendorf Lobind tubes and sonicated for 3 cycles consisting of 5 s of active sonication at 30% amplitude, followed by 15 s on ice. Samples were then centrifuged for 5 min at 15,000 x *g* and the supernatant transferred to a new tube. Protein concentration was determined by bicinchoninic acid (BCA) assay (Pierce). For protein digestion, 100 μg of each sample was aliquoted and volumes normalized with additional lysis buffer. Samples were reduced with 1 mM dithiothreitol (DTT) at room temperature for 30 min, followed by 5 mM iodoacetamide (IAA) alkylation in the dark for another 30 min. Lysyl endopeptidase (Wako) at 1:100 (w/w) was added, and digestion allowed to proceed overnight. Samples were then 7-fold diluted with 50 mM ammonium bicarbonate. Trypsin (Promega) was then added at 1:50 (w/w) and digestion was carried out for another 16 h. The peptide solutions were acidified to a final concentration of 1% (vol/vol) formic acid (FA) and 0.1% (vol/vol) trifluoroacetic acid (TFA) and desalted with a 30 mg HLB column (Oasis). Each HLB column was first rinsed with 1 mL of methanol, washed with 1 mL 50% (vol/vol) acetonitrile (ACN), and equilibrated with 2×1 mL 0.1% (vol/vol) TFA. The samples were then loaded onto the column and washed with 2×1 mL 0.1% (vol/vol) TFA. Elution was performed with 2 volumes of 0.5 mL 50% (vol/vol) ACN. An equal amount of peptide from each sample was aliquoted and pooled as the global pooled internal standard (GIS), which was split and labeled in each TMT batch as described below.

### Isobaric Tandem Mass Tag (TMT) Peptide Labeling

The 610 ROSMAP cases included were analyzed in two separate sets, referred to as Set1 (*n*=400) and Set2 (*n*=210) throughout the Methods. Within each set, cases were randomized prior to TMT labeling by select covariates (i.e., age, sex, PMI, diagnosis) into batches. For Set1 (50 batches), peptides from each individual case and the GIS pooled standards were labeled using the TMT 10-plex kit (ThermoFisher 90406). For Set2 (14 batches), peptides from each individual case and the GIS pooled standards were labeled using the TMTpro 16-plex kit (ThermoFisher 44520). Each batch in Set1 comprised 2 TMT channels with labeled GIS standards with all other channels reserved for individual brain samples. Each batch in Set2 comprised only 1 TMT channel with a labeled GIS standard. Labeling was performed as previously described [44, 84, 85]. Briefly, each sample (containing 100 μg of peptides) was re-suspended in 100 mM TEAB buffer (100 μL). The TMT labeling reagents (5mg) were equilibrated to room temperature, and anhydrous ACN (256 μL) was added to each reagent channel. Each channel was gently vortexed for 5 min, and then 41 μL from each TMT channel was transferred to the peptide solutions and allowed to incubate for 1 h at room temperature. The reaction was quenched with 5% (vol/vol) hydroxylamine (8 μl) (Pierce). All channels were then combined and dried by SpeedVac (LabConco) to approximately 150 μL and diluted with 1 mL of 0.1% (vol/vol) TFA, then acidified to a final concentration of 1% (vol/vol) FA and 0.1% (vol/vol) TFA. Labeled peptides were desalted with a 200 mg C18 Sep-Pak column (Waters). Each Sep-Pak column was activated with 3 mL of methanol, washed with 3 mL of 50% (vol/vol) ACN, and equilibrated with 2×3 mL of 0.1% TFA. The samples were then loaded and each column was washed with 2×3 mL 0.1% (vol/vol) TFA, followed by 2 mL of 1% (vol/vol) FA. Elution was performed with 2 volumes of 1.5 mL 50% (vol/vol) ACN. The eluates were then dried to completeness using a SpeedVac.

### High-pH Off-line Fractionation

High pH fractionation was performed essentially as described [84, 86] with slight modification. Dried samples were re-suspended in high pH loading buffer (0.07% vol/vol NH4OH, 0.045% vol/vol FA, 2% vol/vol ACN) and loaded onto an Agilent ZORBAX 300 Extend-C18 column (2.1mm x 150 mm with 3.5 μm beads). An Agilent 1100 HPLC system was used to carry out the fractionation. Solvent A consisted of 0.0175% (vol/vol) NH4OH, 0.01125% (vol/vol) FA, and 2% (vol/vol) ACN; solvent B consisted of 0.0175% (vol/vol) NH4OH, 0.01125% (vol/vol) FA, and 90% (vol/vol) ACN. The sample elution was performed over a 58.6 min gradient with a flow rate of 0.4 mL/min. The gradient consisted of 100% solvent A for 2 min, then 0% to 12% solvent B over 6 min, then 12% to 40 % over 28 min, then 40% to 44% over 4 min, then 44% to 60% over 5 min, and then held constant at 60% solvent B for 13.6 min. A total of 96 individual equal volume fractions were collected across the gradient and subsequently pooled by concatenation [86] into 24 fractions for Set1 and 48 fractions for Set2. The fractions were then dried to completeness using a SpeedVac.

### Mass Spectrometry Analysis

For Set1, all fractions were resuspended in an equal volume of loading buffer (0.1% FA, 0.03% TFA, 1% ACN) and analyzed by liquid chromatography coupled to tandem mass spectrometry essentially as described [9], with slight modifications. Peptide eluents were separated on a self-packed C18 (1.9 μm, Dr. Maisch, Germany) fused silica column (25 cm × 75 μM internal diameter (ID); New Objective, Woburn, MA) by a Dionex UltiMate 3000 RSLCnano liquid chromatography system (ThermoFisher Scientific). Elution was performed over a 180 min gradient with flow rate at 225 nL/min. The gradient was from 3% to 7% buffer B over 5 min, then 7% to 30% over 140 min, then 30% to 60% over 5 min, then 60% to 99% over 2 min, then held constant at 99% solvent B for 8 min, and then back to 1% B for an additional 20 min to equilibrate the column. Buffer A was water with 0.1% (vol/vol) formic acid, and buffer B was 80% (vol/vol) acetonitrile in water with 0.1% (vol/vol) formic acid. Peptides were monitored on an Orbitrap Fusion mass spectrometer (ThermoFisher Scientific). The mass spectrometer was set to acquire in data dependent mode using the top speed workflow with a cycle time of 3 seconds. Each cycle consisted of 1 full scan followed by as many MS/MS (MS2) scans that could fit within the time window. The full scan (MS1) was performed with an m/z range of 350-1500 at 120,000 resolution (at 200 m/z) with AGC set at 4×10^5^ and maximum injection time 50 ms. The most intense ions were selected for higher energy collision-induced dissociation (HCD) at 38% collision energy with an isolation of 0.7 m/z, a resolution of 30,000, an AGC setting of 5×10^4^, and a maximum injection time of 100 ms. Five of the 50 TMT batches were run on the Orbitrap Fusion mass spectrometer using the SPS-MS3 method as previously described [84]. All higher energy collision-induced dissociation (HCD) MS/MS spectra were acquired at a resolution of 60,000 (1.6 m/z isolation width, 35% collision energy, 5×10^4^ AGC target, 50 ms maximum ion time). Dynamic exclusion was set to exclude previously sequenced peaks for 20 seconds within a 10-ppm isolation window.

For Set2, all fractions were resuspended in an equal volume of loading buffer (0.1% FA, 0.03% TFA, 1% ACN) and analyzed by liquid chromatography coupled to tandem mass spectrometry essentially as described [9], with slight modifications. Peptide eluents were separated on a self-packed C18 (1.9 μm, Dr. Maisch, Germany) fused silica column (15 cm × 75 μM internal diameter (ID); New Objective, Woburn, MA) by an EASY-nLC 1200 liquid chromatography system (ThermoFisher Scientific). Elution was performed over a 45 min gradient with flow rate at 400 nL/min. The gradient was from 5% to 35% over 37 min, then 35% to 99% over 1 min, then held constant at 99% solvent B for 8 min, and then back to 1% B for an additional 7 min to equilibrate the column. Buffer A was water with 0.1% (vol/vol) formic acid, and buffer B was 80% (vol/vol) acetonitrile in water with 0.1% (vol/vol) formic acid. Peptides were monitored on a Q-Exactive HFX mass spectrometer (ThermoFisher Scientific). The full scan (MS1) was performed with an m/z range of 410-1600 at 120,000 resolution (at 200 m/z) with AGC set at 3×10^6^ and maximum injection time 50 ms. The top 20 most intense ions were selected for higher energy collision-induced dissociation (HCD) at 32% collision energy with an isolation of 0.7 m/z, a resolution of 45,000, an AGC setting of 2×10^5^, and a maximum injection time of 96 ms. Dynamic exclusion was set to exclude previously sequenced peaks for 20 seconds within a 10-ppm isolation window.

### Database Searches and Protein Quantification

All RAW files (1,200 RAW files from TMT-MS analysis of ROSMAP Set1 and 672 RAW files from TMT-MS of ROSMAP Set2) were analyzed using the Proteome Discoverer suite (version 2.4, ThermoFisher Scientific). MS2 spectra were searched against the UniProtKB human proteome database containing Swiss-Prot human reference protein sequences (20338 target proteins). The Sequest HT search engine was used and parameters were specified as follows: fully tryptic specificity, maximum of two missed cleavages, minimum peptide length of 6, fixed modifications for TMT tags on lysine residues and peptide N-termini (+229.162932 Da) and carbamidomethylation of cysteine residues (+57.02146 Da), variable modifications for oxidation of methionine residues (+15.99492 Da) and deamidation of asparagine and glutamine (+0.984 Da), precursor mass tolerance of 20 ppm, and a fragment mass tolerance of 0.05 Da for MS2 spectra collected in the Orbitrap (0.5 Da for the MS2 from the SPS-MS3 batches).

Percolator was used to filter peptide spectral matches (PSMs) and peptides to a false discovery rate (FDR) of less than 1%. Following spectral assignment, peptides were assembled into proteins and were further filtered based on the combined probabilities of their constituent peptides to a final FDR of 1%. A multi-consensus was performed to achieve parsimonious protein grouping across individual batches and both sets of ROSMAP samples. In cases of redundancy, shared peptides were assigned to the protein sequence in adherence with the principles of parsimony. As default, the top matching protein or “master protein” is the protein with the largest number of unique peptides and with the smallest value in the percent peptide coverage (i.e., the longest protein). Reporter ions were quantified from MS2 or MS3 scans using an integration tolerance of 20 ppm with the most confident centroid setting. Only unique and razor (i.e., parsimonious) peptides were considered for quantification.

### Controlling for Batch-specific Variance Across Proteomics Datasets

A tunable median polish approach (TAMPOR) was used to remove technical batch variance in the proteomic data, as previously described [31, 32]. Following a multi-consensus database search and protein quantification across two sets of ROSMAP tissues, batch effects in the first set (set1) of ROSMAP samples (50 TMT-10 plex batches) and the second set (set2) of ROSMAP samples (14 TMT-16 plex batches) were normalized iteratively in two steps essentially as described [32]. After removal of intra-set batch effects in set1 and set2 separately, all samples except set-specific GIS samples were processed jointly with TAMPOR into a single reassembled consensus sample–protein matrix using the median of within-cohort pathology free control cases as the central tendency, enforcing that the population of all log_2_(ratio) output for control samples within the final 610 ROSMAP samples would tend toward 0. Additional details on the data input and TAMPOR output can be found on https://www.synapse.org/#!Synapse:syn32280722.

### Regression of Covariates

The ROSMAP cases were subjected to nonparametric bootstrap regression by subtracting the trait of interest (age at death, sex, or PMI) times the median estimated coefficient from 1000 iterations of fitting for each protein in the log_2_(abundance) matrix as previously described [32]. Ages at death used for regression were uncensored.

### Sample Network Generation with MONET M1 Algorithm

The three top-performing methods from the DMI DREAM Challenge were compiled in the MONET toolbox and released to the public for use (https://github.com/BergmannLab/MONET.git) [40]. We selected the M1 method from this toolbox to build a sample-wise network. The M1 method uses the Girvan-Newman modularity optimization method to group like features into modules [87]. MONET M1 has expanded on the traditional modularity optimization to a multiresolution approach by searching the network at multiple topological scales. The authors have added the resistance parameter, *r*, which averts genes from joining modules. If *r* = 0 the method returns to Newman and Girvan’s original modularity optimization; *r* > 0 reveals network substructure; and *r* < 0 network superstructure [88]. The parameter *r*, is fit to four user-provided hyperparameters: minimum module size, maximum module size, desired average degree, and desired average degree tolerance to produce a network described by the parameters.

After batch-correction and regression as described above, *n* = 610 cases were split into two groups defined by the ROSMAP cognitive diagnosis score: cognitively impaired cases (*n* = 377, *cogdx* ∈ {2,3,4,5}) and non-cognitively impaired/other cases (*n* = 233, *cogdx* ∈ {1,6}). An expression data matrix of the cognitively impaired cases *n* = 377 and *n* = 7,723 *log*_2_ (*protein abundance*) was created (proteins with greater 50% missingness in the 377 cases were removed). The adjacency function in WeiGhted Correlation Network Analysis (WGCNA) was used to build the adjacency matrix with parameters: soft threshold power = 8, type=“signed”, corFnc=“bicor”, and the corOptions parameter set to use pairwise complete correlation [89]. The soft threshold power was determined using scale free topology analysis based on the following two guidelines: 1) The power in a plot of power (x) vs R^2^ (y) should be where the R^2^ has approached an asymptote, usually near or above 0.80, and 2) the mean and median connectivity at that power should not be exceedingly high, preferably around 100 or less. The power at which these criteria are met is a tradeoff between removing correlations due to chance and maintaining as many correlations in the data as possible for the clustering algorithm to distinguish modules. As M1 takes an edge list as input, the adjacency upper triangle correlation values were used to populate the weights of unique pairwise correlations in the edge list. No sparsification of the edge list was applied.

The hyperparameters were optimized using a grid search by varying minimum module size, *i* ∈ {5, 10, 25, 50}, maximum module size, *j* ∈ {100, 150, 200, 250, 300, 350, 376}, and desired average degree, *k* ∈ {25, 50}. The desired average degree tolerance was left at the default value of 0.2. The optimal parameter set was defined as the set that minimized the percentage of cases not assigned to a module and the maximum module size (**Fig. S1, Table S1**). The final parameters selected were *i* = 5, *j* = 200, *k* = 50, which built a network with 3 modules and 5.31% (*n* = 20) cases not assigned to a module. This final network was used in to define the three subtypes of AD observed in the ROSMAP cohort.

Using the expression data matrix and the modules assignment list, module eigenvectors were defined, using the moduleEigengenes function in WGCNA. The eigenvector is the module’s first principal component and explains covariance of all cases within each module [citation]. Using the signedKME function in WGCNA, a table of bicor correlations between each case and each module eigenvector was obtained; this module membership measure is defined as kME. Additional details of the data input and MONET M1 output can be found on https://www.synapse.org/#!Synapse:syn32567811.

### MONET M1 Bootstrap Validation

To validate the robustness of the clustering, we performed 100 rounds of bootstrapped reclustering, by withhold 20% of the samples and employing the MONET M1 method on the subset (**Fig. S2, Table S2**). Due to the decrease in sample number, the hyperparameters were adjusted to *i* = 5, *j* = 160, *k* = 50, where the maximum module size was decreased by 20%. If there are truly unique molecular subtypes, then repeated clustering should produce similar sets of clustered samples. To assess the probability of sample pairs clustering together, we calculated the rate at which each pair is assigned to them same cluster (termed the paired percentage). The average paired percentage per cluster and variance of the percentage were tracked for the 100 bootstrap steps to ensure the bootstrapping was well converged. A heatmap of the paired percentage shows that the clustering is stable and repeated rounds of MONET M1 reproduces the three observed subtypes. A hypergeometric Fisher’s exact test (FET) was performed on each of the bootstrap steps against the original MONET M1 cluster assignments and showed class-specific overlap. Along with the rate that sample pairs clustered together, the reproducibility was also assessed using the module preservation function in WGCNA. *Z*_summary_ composite preservation scores were obtained using the MONET M1 network (n=377) as the template versus each bootstrap step, with 100 permutations. Random seed was set to 1 for reproducibility, and the quickCor option was set to 0. The summary z scores were then averaged across the 100 bootstrap steps and the standard deviation was calculated. All three subtypes were preserved across the 100 bootstrap replicates (above the blue line *q*=0.05).

### Uniform Manifold Approximation and Projection (UMAP) Dimension Reduction

Supervised dimensionality reduction was performed using UMAP (umap-learn v0.5.2) in Python (v3.9) with the following settings: n_neighbors = 10, n_components = 2, metric = Euclidean, and min_dist=1. A supervised UMAP embedding was generated for the 357 cases in the MONET M1 classes using the n=7,723 *log_2_(protein abundance)* as features and the three MONET M1 classes as target labels. The three resultant clusters mirror those of MONET M1 network analysis with only 1 case out of the 357 assigned to a cluster different from its original class.

### Biological Organization of Protein Expression Matrix

To highlight the biological trends between the three clusters produced by MONET M1, the proteins used as features in the network (*n* = 7,723) were grouped into the protein modules generated in our deepest AD consensus network [32]. For each protein module, proteins were sorted by kME. Protein modules were then organized into relatedness order determined by previous WGCNA analysis. This highlighted the systems-based divergences and biological trends between MONET M1 classes allowing us to character the classes by their proteomic expression profiles. For visualization, samples within the 3 MONET M1 classes were also sorted by their class specific kME (**Table S2**). Therefore, the first sample is the “eigensample” for the MONET M1 class.

### ROC Analyses

For each protein, a support vector machine classifier was used to classify individuals by class using one-vs-rest strategy. Due to the multi-class nature of the model, each class identifier was binarized prior to training. To choose the model for which predictions are made, the one-vs-rest heuristically chooses the binary classifier with the highest confidence. After a model was created, the receiver operating characteristic (ROC) metric was used to evaluate each classifier. The area under the curve (AUC) was calculated for each peptide in each classifier. The peptides were then sorted based on the AUC values and the top ten were plotted for each class.

### Protein Correlation to Traits

To assess which proteins most strongly correlated to clinicopathological traits, the WGCNA R netScreen function was used. All 7,723 proteins were correlated to the provided ROSMAP across the 357 samples (**Table S8**). The top 10 positive and top 10 negatively correlated proteins were manually filtered from the analysis. The z-score for these proteins were plotted for each of the three Classes and the NCI group to visualize the expression differences of these proteins across the unique subtypes.

### Other Statistics

Statistical analyses were performed in Python v3.7 and visualized using matplotlib package v3.5.1. Correlations were performed using the biweight midcorrelation function as implemented in the WGCNA R package (**Tables S4 and S8**). Boxplots represent the median, 25^th^, and 75^th^ percentile extrema, the edges of a box represent the interquartile of the data within a group. Whiskers are drawn at the maximum and minimum value of the data set. Points greater than 1.5 times the interquartile range are classified as outliers. Comparisons between two groups were performed by *t* test. Comparisons among three or more groups were performed with one-way ANOVA with Tukey or Bonferroni pairwise comparison of significance (**Tables S3, S5, S6, S9**). *P* values were adjusted for multiple comparisons by false discovery rate (FDR) correction where indicated.

## Author Contributions

Conceptualization, L.H., E.K.C., E.B.D., E.C.B.J., J.J.L., A.I.L., and N.T.S.; Methodology, L.H., E.K.C., E.B.D., R.U.H., D.M.D., and N.T.S.; Investigation, L.H., E.K.C, E.B.D., D.M.D., L.Y., and N.T.S.; Formal Analysis, E.K.C., E.B.D., R.U.H.; Writing – Original Draft, L.H., E.K.C., E.B.D., and N.T.S.; Writing – Review & Editing, L.H., E.C.B.J., J.J.L., A.I.L., and N.T.S.; Funding Acquisition, A.I.L. and N.T.S.; Resources, P.L.D. and D.A.B.; Supervision, A.I.L., J.J.L., and N.T.S.

## Data and Code Availability

Raw mass spectrometry data from the ROSMAP dorsolateral prefrontal cortex tissues can be found at https://www.synapse.org/#!Synapse:syn17015098. Pre- and post-processed protein expression data and case traits related to this manuscript are available at https://www.synapse.org/ADsubtype. The algorithm used for batch correction is fully documented and available as an R function, which can be downloaded from https://github.com/edammer/TAMPOR. The results published here are in whole or in part based on data obtained from the AMP-AD Knowledge Portal (https://adknowledgeportal.synapse.org). The AMP-AD Knowledge Portal is a platform for accessing data, analyses, and tools generated by the AMP-AD Target Discovery Program and other programs supported by the National Institute on Aging to enable open-science practices and accelerate translational learning. The data, analyses, and tools are shared early in the research cycle without a publication embargo on secondary use. Data are available for general research use according to the following requirements for data access and data attribution (https://adknowledgeportal.synapse.org/#/DataAccess/Instructions). Additional ROSMAP resources can be requested at www.radc.rush.edu.

